# Guitarfishes are plucked: undermanaged in global fisheries despite declining populations and high volume of unreported international trade

**DOI:** 10.1101/2022.10.05.510982

**Authors:** C. Samantha Sherman, Colin A. Simpfendorfer, Alifa B. Haque, Eric D. Digel, Patrick Zubick, Jonathan Eged, Jay H. Matsushiba, Glenn Sant, Nicholas K. Dulvy

## Abstract

Some sharks and rays are subject to fisheries catch and international trade regulations. However, the Guitarfishes (family Rhinobatidae) are a highly threatened group with minimal regulations. Substantial underreporting of catch and broad commodity codes for traded products are masking the true volume of Guitarfishes included in international trade. Here, we collate international trade information for Guitarfishes that have not readily been documented in trade, possibly due to poor resolution of molecular genetic markers, to begin to document the extent of trade. We assess the shortfall in fisheries management (M-Risk) for all species of Guitarfish based on 99 assessments across 28 countries. Globally, Guitarfishes are inadequately managed, with an average M-Risk of 45% of an ideal score, resulting in 76% of species being threatened globally. The high and unregulated catch and trade volume, paired with the management shortfalls, require global integrated improvement in fisheries management, supported by regulating international trade to sustainable levels.

## 1 Introduction

The rhino rays are shark-like rays of the Rhinopristiformes Order and is comprised of Sawfishes (family Pristidae), Wedgefishes (family Rhinidae), Giant Guitarfishes (family Glaucostegidae), Guitarfishes (family Rhinobatidae), and Banjo Rays (family Trygonorhinidae)(Dulvy and Simpfendorfer 2022). Rhino rays are amongst the most imperilled of elasmobranchs, with 79% (46 of 58 species) of data-sufficient species listed in an IUCN Red List threatened category (Dulvy et al. 2021, IUCN 2022). Aside from the Banjo Rays, which have quite different life history characteristics from the rest of the rhino rays, they are highly valued for their fins and snouts, and with their meat considered to be of good quality (Haque et al. 2021). As a result, catches over recent decades have increased, and many regularly enter the international trade (Moore 2017, Cardeñosa et al. 2020, Choy et al. 2022). The fins of Sawfishes, Wedgefishes and Giant Guitarfishes are the most highly prized and highly valued fins and are known as “Qun Chi” and these fins are worth USD$185–964 per kg (Hau et al. 2018, Jabado 2019). As a result, the three families with the largest body sizes and greatest proportions of threatened species (Sawfishes, Giant Guitarfishes, and Wedgefishes) were added to Appendix II of the Convention on International Trade in Endangered Species (CITES) in 2007 (Sawfishes) and 2019 (Giant Guitarfishes and Wedgefishes) to ensure legal and sustainable international trade (Dent and Clarke 2015, Dulvy et al. 2021). The Guitarfish family consists of 37 species in three genera (*Acroteriobatus*, 10 spp.; *Pseudobatos*, 9 spp.; *Rhinobatos*, 18 spp.) that occur in tropical coastal waters globally (Last et al. 2016). The trade in Guitarfishes remains data-poor and unregulated.

Guitarfishes are caught in a wide range of gears (especially trawls, gillnets and longlines) in both large-scale and small-scale artisanal and subsistence fisheries throughout their ranges (Choy et al. 2022, Seidu et al. 2022b). Guitarfishes are targeted and are increasingly being retained as incidental catch to be used for their skins for leather, and fins, meat, and rostra as food for human consumption (Jabado et al. 2021b). Trade of Guitarfishes, and the drivers of their trade, are poorly documented and sits within the broader international trade of sharks and rays where underreporting is common. For example, in Indonesia, there was an estimated underreporting of over $60 million dollars in shark fin and meat from 2012–2018 (Prasetyo et al. 2021). The Guitarfishes tend to have smaller less valuable fins than the larger rhino rays (Temple 2018). However, as the larger Sawfishes, Wedgefishes, and Giant Guitarfishes have been depleted there is increasing evidence that the smaller species are being targeted and retained in the belief that they will also garner the high values of the larger rhino rays (Seidu et al. 2022b). As Guitarfishes are primarily distributed through countries with higher levels of artisanal fishing and lower capacity to manage catch, there is concern for the persistence of Guitarfishes in these heavily fished regions. Guitarfishes have low productivity. For example, two *Pseudobatos* species (*P. horkelii* Brazilian Guitarfish and *P. productus* Shovelnose Guitarfish) have lower intrinsic rates of population increase when compared to seven other species of rhino rays (D’Alberto et al. 2019). This indicates these species have higher intrinsic susceptibility to being overfished than rhino ray species that are already subject to increased management through international legislation, including being listed on CITES Appendix II (Pardo et al. 2016, Fernando et al. 2022).

Here, we assess the efficacy of fisheries management for all Guitarfish species using a rapid management risk assessment - M Risk (Sherman et al. 2022b). Specifically, we consider three questions about the management of Guitarfishes: (1) are Guitarfish adequately managed throughout their geographic range to ensure their ongoing survival? (2) which species have the best management and which have the most limited management?, and (3) which aspects of management are most prevalent and lacking across countries and species?

## 2 Methods

### 2.1 Selection of Management Units

The methods here follow closely a similar study on the Requiem Sharks (Family Carcharhinidae) and the complete method is described in detail in Sherman (*et al*. 2022). Management units were selected based on their proportional contribution to global shark and ray catch (2010-2019, inclusive) (FAO 2019). We included the 21 countries that cumulatively report >80% of the global Chondrichthyan catch in addition to nine countries with the highest catch in each FAO fishing region to ensure global representation (total = 30). Assessments were completed for each species of Guitarfish that occurred in any of the management units considered. Guitarfish did not occur in all 30 countries considered. Therefore, the final number of management units that included at least one species of Guitarfish, and included in our analyses, was 28.

### 2.2 M-Risk: Management Risk Assessment

Species assessments were completed by scoring management risk against 21 measurable attributes, as documented in Sherman *et al*. (2022). The 21 attributes were split into four classes, three scored for all management units (Management System [*n*=5 Attributes], Fishing Practises & Catch [*n*=5], Compliance, Monitoring, & Enforcement [*n*=5]), and two that were specific to either a country (*n*=4) or RFMO (*n*=2) (**Supplementary Information**). As the catch of Guitarfishes did not occur in any RFMO, only 19 of the M-Risk attributes were considered for this analysis.

### 2.3 Scoring of Attributes

All attributes were scored in an ordinal manner with a narrow range of scores to ensure consistency (i.e., 0-3, 0-4, to 0-5) such that higher scores indicated management with a higher likelihood of sustainable outcomes (for full details see Sherman *et al*., 2022b). Assessments were completed based on information sourced through exhaustive internet searches and national fisheries department websites. Where English was not the official language, searches were completed in the official language and documents were translated using Google translate. To accommodate the potential for ambiguous translations, points were allocated generously when there was uncertainty in the translated information. However, if no information for an attribute was found, a precautionary score of zero was given. The final ‘Management Score’ was expressed as a percentage of the total possible points. This score indicates progress towards the ideal management, therefore, a score greater than 75% should be considered a well-managed fishery for sharks and rays (Sherman et al. 2022a).

### 2.4 Additional Catch, Trade, and Species Data

Reported catch and trade data were sourced from the Food and Agriculture Organisation (FAO) using their software FishStatJ to search two databases: (i) Global Fishery and Aquaculture Production Statistics (v2022.1.1) and (ii) Global Fish Trade Statistics (v2022.1.0; FAO 2022). Additionally, unreported catch data was downloaded from the Sea Around Us website (www.seaaroundus.org; Pauly et al. 2021). Google scholar searches were completed to find additional published data and evidence for Guitarfish catch and trade. Evidence was found for domestic catch, export, and import of Guitarfishes for 75 countries. Therefore, a zero value does not indicate that a country does not catch, export, or import Guitarfishes, rather no data was found to say otherwise.

Threat level was determined by searching the IUCN Red List of Threatened Species website, where species distribution maps were also sourced from (www.iucnredlist.org). These were sourced after the 2021-03 update, which was the final update from the Global Shark Trends reassessment of all 1,199 species as summarised here (Dulvy et al. 2021).

## 3 Results

### 3.1 What is the scale of catch and trade of Guitarfishes?

Guitarfishes (family Rhinobatidae) are caught widely throughout the tropics and subtropics. Global reported catch of Guitarfishes are limited as the FAO databases only reported to the family (Rhinobatidae) or one of three species (*Pseudobatos percellens* Chola Guitarfish, *R. planiceps* Pacific Guitarfish, and *R. rhinobatos* common Guitarfish; FAO, 2022; **Supplementary Data S1**). Nonetheless, reporting for these four groups has averaged 3,914.14 mt/yr over the past ten years from 18 different countries (FAO 2022). However, there is no catch of Guitarfishes reported to the FAO from major fishing nations which are range states of Guitarfishes (such as India, Singapore, and Tanzania, among others) where Guitarfish catches have been documented in local fish markets (Barrowclift et al. 2017, Bineesh et al. 2017, Wainwright et al. 2018). Reconstructed catch data from the Sea Around Us project shows the peak of global Guitarfish catches occurred in the early 1980s, followed by another increase in catch in 2010. However, the major Guitarfish catching countries during the first peak differed as Peru and Brazil dominated the catches during the peak in the 1980s and Mexico and Sierra Leone significantly increased their contribution to global Guitarfish catch in 2010 (Pauly et al. 2021). In Brazil, where catch of Brazilian Guitarfish is prohibited, there are still products sold in markets as Guitarfish, much of which is mislabelled (Bernardo et al. 2020). Guitarfishes are retained in many countries for their meat, which is consumed fresh or dried and preserved (Márquez-Farías 2007, Jabado et al. 2021b). In Tunisia, skins of Guitarfishes are used locally to make drums (Newell 2017). In Mexico, shovelnose Guitarfish accounts for 93% of ray catch in the Pacific, and 45% of ray catch in the artisanal Gulf of California ray fishery (Márquez-Farías 2007). In Pakistan, there are an estimated 5,700 vessels using benthic gear that catch and retain rhino rays, which are sold at local markets (Moazzam and Osmany 2020). In total, we found evidence (both landings report and anecdotally through published papers) that Guitarfishes are caught in at least 72 countries across all six inhabited continents (**Figure 1A; Supplementary Data S1**).

**Figure 1.**
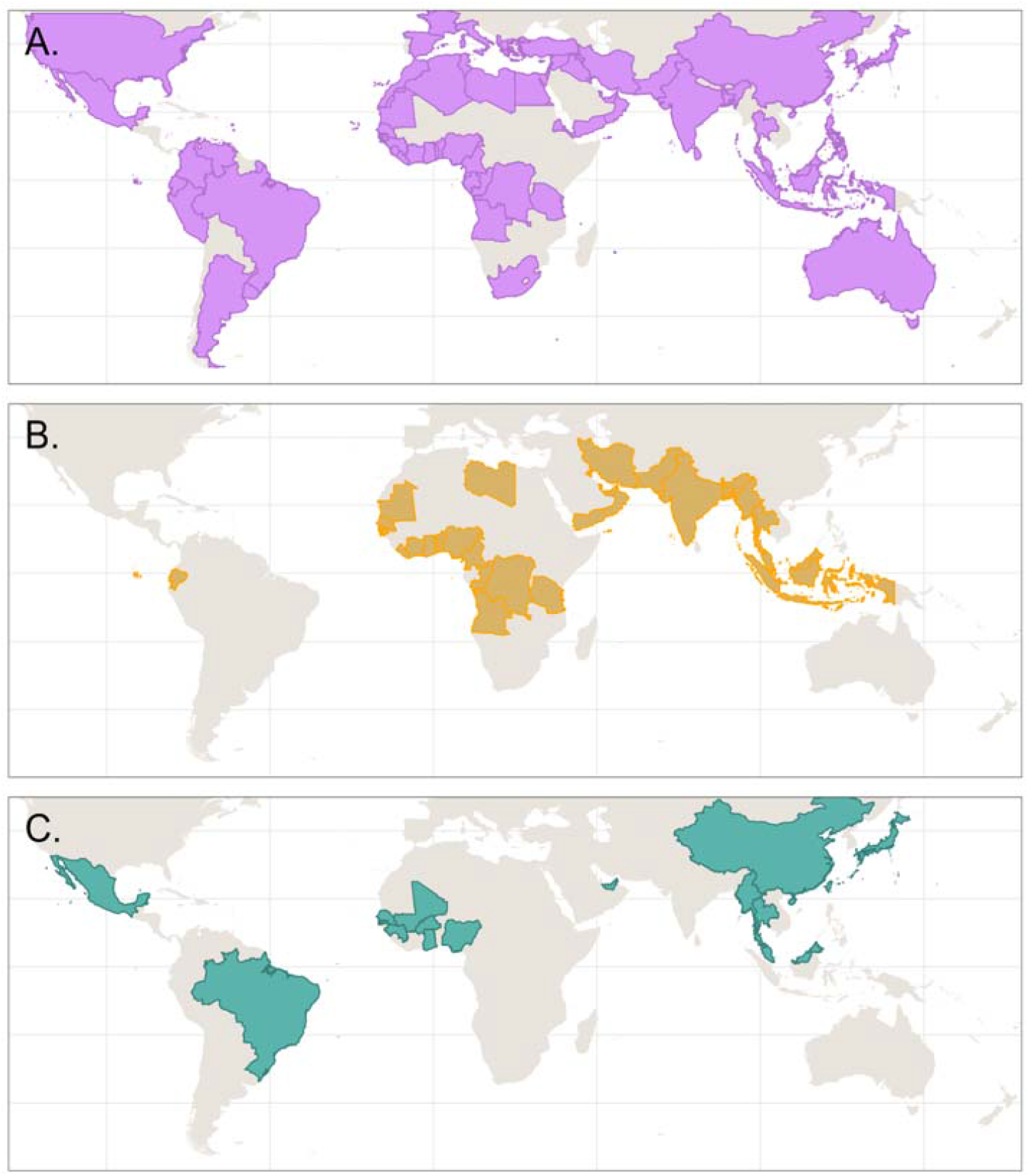
Global maps of (A) countries with documented catch of guitarfishes within their Exclusive Economic Zone (EEZ), (B) countries with documented export of Guitarfishes (including middleman countries), and (C) countries with documented import of Guitarfishes. Data sources can be found in **Supplementary Data S1**. Non-coloured countries do not indicate non-involvement in catch and trade, rather no available data.

There is no available estimate for the volume of international trade of Guitarfishes. Non-standardized commodity codes mask catch and trade details, making the full trade of this group impossible to determine (FAO 2022). Their fins are in demand and fetch high prices on the international market, and are exported from Bangladesh (if large enough), Indonesia, and several African countries to China (Haque and Spaet 2021, Jabado et al. 2021b, Prasetyo et al. 2021). In Ghana, while meat is sold for local consumption, fins from Guitarfishes are exported either through middlemen or directly from the fishers to buyers in The Gambia, Mali, and Senegal (Seidu et al. 2022b). Skins of Guitarfishes are used to produce a luxury leather and are exported from Bangladesh to Myanmar (Haque et al. 2018). Almost all of these exported products are unreported, despite some exporters claiming to have traded up to 600 tonnes of shark and ray product annually (Haque and Spaet 2021). Some species of Guitarfishes or juveniles, are too small to fetch high prices in international trade, however, they are often consumed locally, or other products may be traded (i.e., skins used as a luxury leather from *R. schlegelii*, brown Guitarfish)(Rigby et al. 2021). There is likely a high volume of Guitarfish fin export coming from Indonesia, where international trade is routed through Jakarta and Surabaya before being exported to Singapore, Thailand, Malaysia, and China (SEAFDEC 2006). Guitarfishes are also landed directly at Singaporean fish markets (Wainwright et al. 2018). However, fishermen at these markets estimate >65% of their fishing occurs in Indonesian and Malaysian waters and just 2% within the Singapore EEZ (Clark-Shen et al. 2021). These examples indicate there is a high volume of international trade of Guitarfishes that is originating from at least 33 countries across four continents, exported to at least 17 countries, and mostly undocumented and unreported (**Figure 1B,C**; **Supplementary Data S1**). Due to the small geographic range of most Guitarfish species (few span multiple continents), this also proves there are many species included in the international trade of Guitarfishes.

### 3.2 Are Guitarfishes adequately managed across their geographic range?

Guitarfishes are not adequately managed by fisheries globally despite their global distribution with hotspots in the Eastern Central Atlantic Ocean, the Eastern Central Pacific Ocean, the Northwestern Indian Ocean as well as South Africa to Tanzania (**Figure 2A; Supplementary Data S2**). Across 99 management assessments of 33 Guitarfish species from 28 countries spanning all six inhabited continents, the average management risk was high, with an average of only 44.6% (± 1.3, n = 99) of ideal management in place. When compared to all other assessed Chondrichthyan families, the three rhino ray families had the three lowest average management scores globally, indicating the fewest regulations to their catch and management (Sherman et al., unpublished data). Regionally, the management shortfall for Guitarfishes was greatest in Africa (mean management score 36.0% ± 3.5, n = 20) and Asia (38.9% ± 1.3, n = 38; **Figure 2B,C**). Both continents were much lower than the remaining regions: South America (52.7 ± 1.3, n = 14), North America (53.3% ± 2.2, n = 21), Oceania (55.3% ± 20.2, n = 2), and Europe (62.5% ± 2.8, n = 4; **Figure 2B,C**).

**Figure 2.**
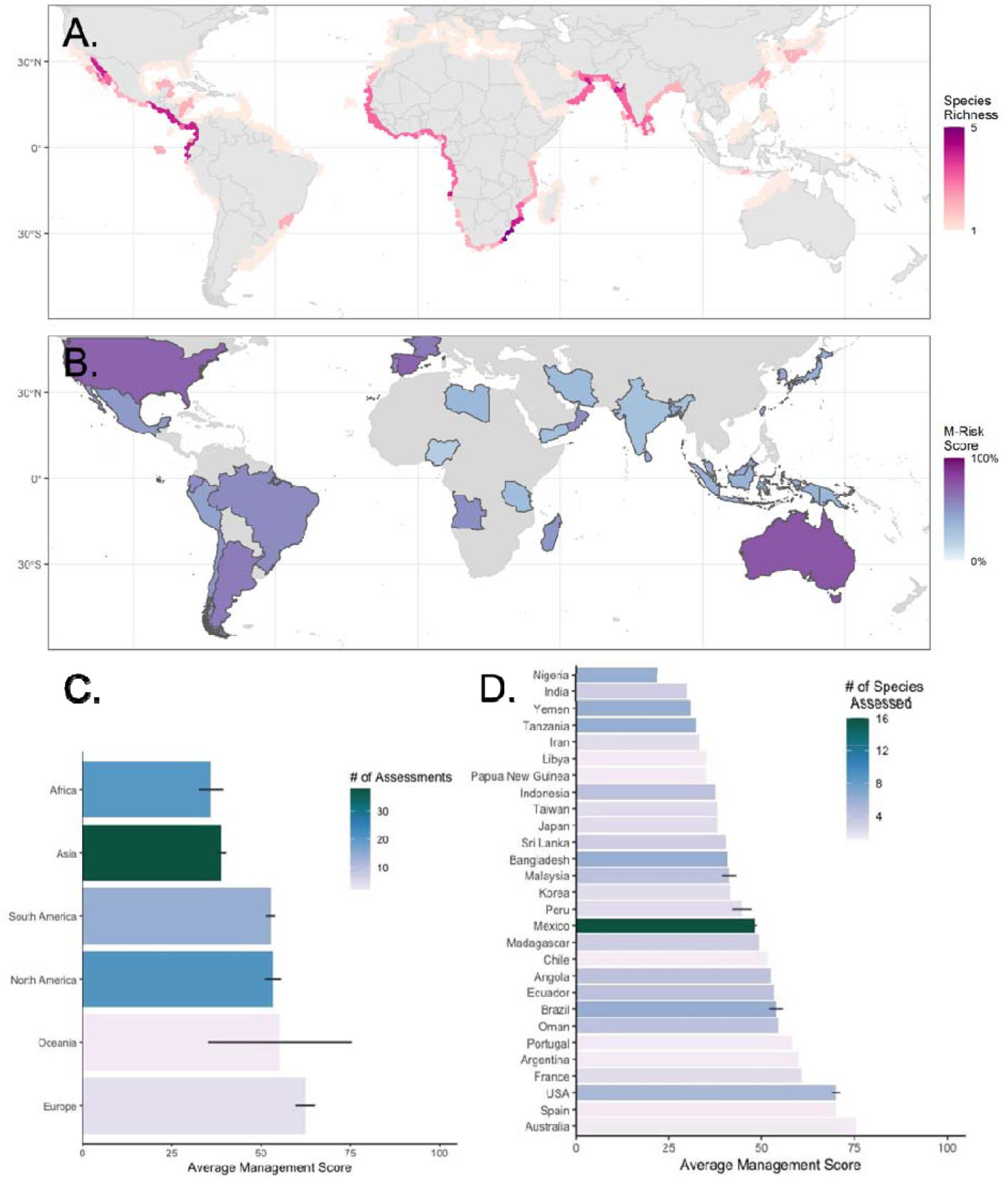
(*This figure should be colour). Global maps of (A) species richness of 34 guitarfish* (family Rhinobatidae; full species list in **Supplementary Data S1**), (B) average management scores for guitarfish in each country assessed. Average management score across different (C) continents, (D) countries. Error bars indicate 95% confidence intervals. *Three species were not mapped as they were recently described and their full range is unknown (details in **Supplementary Data S1**).

The majority of countries considered (60.7%, n = 17 of 28 countries) had average management scores for Guitarfishes lower than half (50%) of an ideal score. The countries with the lowest scores included those among the top shark and ray catching countries in the globally (Okes and Sant 2019). The three lowest scoring countries were Nigeria (21.9% ± 0.0, n = 6), India, (30.0% ± 0.0, n = 3), and Yemen (31.0% ± 0.0, n = 6), respectively (**Figure 2B,D**). The countries with the highest scores were Australia (75.4%, n = 1), the USA (70.0% ± 1.1, n = 5), and Spain (70.0%, n = 1). France and Argentina also had scores over 60% (60.8% and 60.0%, respectively), but only one or two assessments were completed in each of those countries, increasing uncertainty. For countries with a higher diversity of Guitarfishes, and therefore more assessments, the highest scoring countries were the USA, Oman (54.4% ± 0.0, n = 4), and Brazil (53.9% ± 2.0, n = 6) (**Figure 2B,D**).

### 3.3 Which species are adequately managed?

Any species that does not have a management score of 100% is lacking in some aspects of their management. By that criterion - across all 99 assessments, all 33 species are undermanaged. The species with the highest management scores overall are Goldeneye Guitarfish (*Rhinobatos sainsburyi:* 75.4%, n = 1) found in northern Australia, the Shovelnose Guitarfish (58.3% ± 6.5, n = 4) found around Baha, Mexico, and Brazilian Guitarfish (58.3% ± 1.4, n = 3) found in Southwest Atlantic (**Figure 3; Supplementary Data S2**). The Brazilian Guitarfish is one of the most threatened Guitarfishes and is listed as Critically Endangered (Pollom et al. 2020). As a result of the collapse of the Brazilian Guitarfish population, their landing was prohibited in Brazilian fisheries, along with other species-specific regulations, like the creation of marine protected areas at aggregation sites (De-Franco et al. 2012, Anderson et al. 2021). Alternatively, the four lowest scoring species were all assessed in just a single country. Three of these species are endemic only to their respective countries. The lowest scoring species, Socotra Blue-Spotted Guitarfish (*Acroteriobatus stehmanni*: 31.0%) was endemic to and only assessed in Yemen. The next two lowest, Zanzibar Guitarfish (*A. zanzibarensis*: 32.5%) and Slender Guitarfish (*R. holcorhynchus*: 32.5%) were both scored only in Tanzania. Finally, the Papuan Guitarfish (*R. manai*: 35.1%) was endemic to and only scored in Papua New Guinea (**Figure 3**). Only the slender Guitarfish is wide ranging from South Africa to Kenya. Therefore, if management does not improve in these countries, it is possible these species will be overexploited, leading to irreversible population declines, if this has not occurred already.

**Figure 3.**
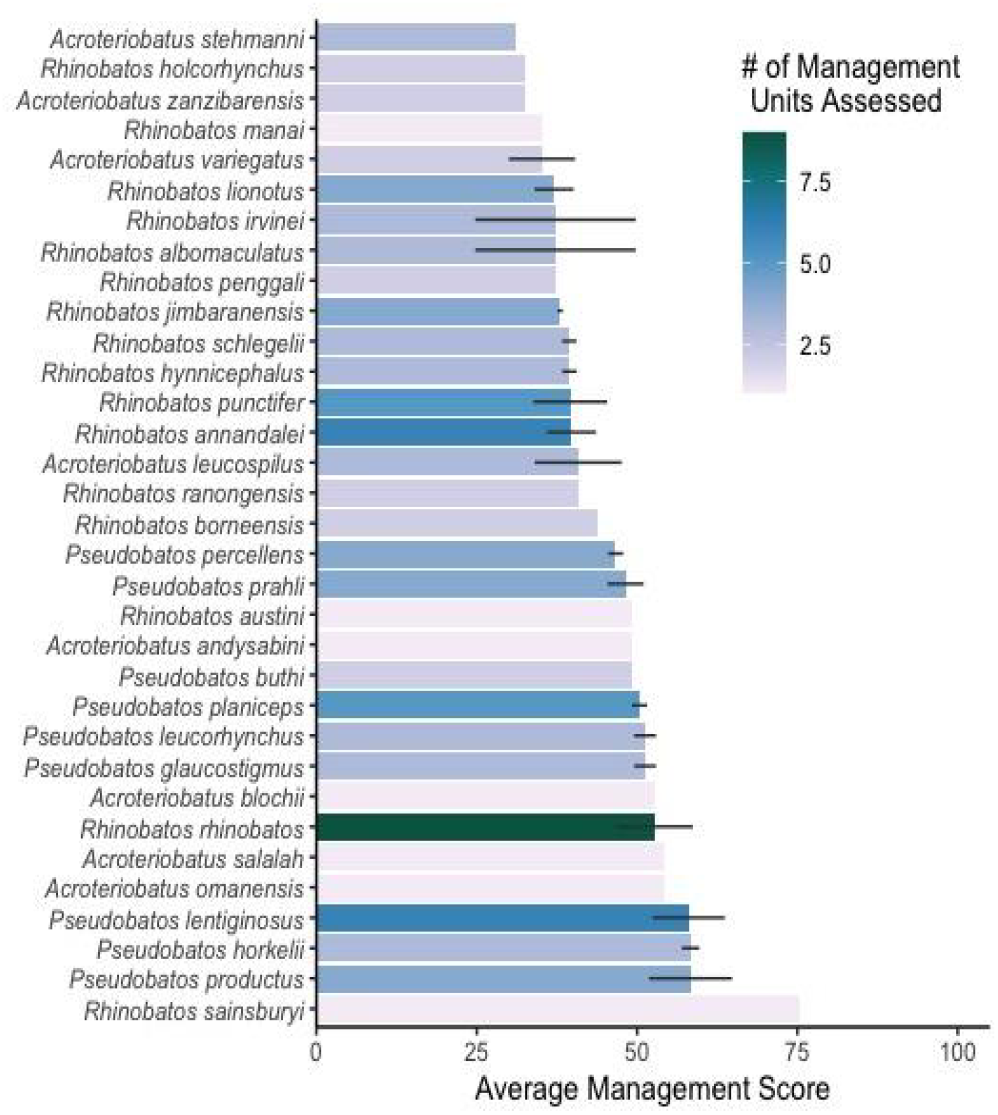
(*This figure should be colour). Average management score for each species assessed. Error bars indicate 95% confidence intervals.

There was a considerable range in management scores for the single highest and lowest scoring Guitarfish assessment spanning 60%. The lowest single score was for three species in the artisanal Nigerian fishery, all scoring 12.3% (Whitespotted Guitarfish, *R. albomaculatus*; Spineback Guitarfish, *R. irvinei*; and common Guitarfish). The highest single score was 75.4% for Goldeneye Shovelnose ray in the Pilbara Trap and Trawl fishery in Western Australia. However, the species with these scores are found in different regions, indicating some species are at much higher risk to undermanagement than others.

Guitarfishes tend to not be as widely distributed as other sharks and rays with an average geographic range of ∼175,000 km^2^ (compared to 14,371,164 km^2^ for Carcharhinidae, approximately 82X larger), meaning national and regional management has a proportionately larger effect on population sizes of Guitarfishes than for species with larger ranges spanning more jurisdictions. The geographic ranges of Guitarfishes will likely become smaller as the taxonomy of species complexes is resolved. For example, two new ‘blue-spotted’ Guitarfishes (Malagasy Blue-Spotted Guitarfish, *A. andysabini* and Socotra Blue-Spotted Guitarfish) were recently described with a revision of Greyspot Guitarfish (*A. leucospilus)* and there is further uncertainty around the status of Zanzibar Guitarfish and it’s relationship or otherwise to Stripenose Guitarfish (*A. variegatus*) found off southern India (Weigmann et al. 2021, Dulvy and Simpfendorer 2022). Additionally, nearly two-thirds (62%, 23 of 37 species) of Guitarfishes are threatened (i.e., Vulnerable, Endangered or Critically Endangered) according to IUCN Categories and Criteria (IUCN Standards; Petitions Subcommittee 2019, Dulvy et al. 2021, IUCN 2022). A further seven species have either not been evaluated or are listed as Data Deficient. If these species are proportionally as threatened as the data sufficient species (77%, 23 of 30), this would increase the proportion of all Guitarfishes threatened to 76% (5.4 of 7 would be threatened; **Supplementary Data S2**)(Walls and Dulvy 2020).

### 3.4 Which aspects of management are most prevalent and lacking

Across all 28 assessed countries, the attributes with the highest average scores were lower resolution ‘structural’ attributes related to general fisheries management, such as: if a regulatory body was present (94.6% ± 3.1), engagement with CITES (92.9% ± 3.1), and management of IUU (81.5% ± 5.5) (**Figure 4A**). Attributes with lower scores were those that required species-specific regulations or knowledge, including: understanding the species’ stock status (7.1% ± 4.0), species-specific compliance measures (14.5% ± 4.9), and taxonomic resolution of landing limits, if any exist (23.8% ± 7.2) (**Figure 4A**). These scores indicate that while most countries have the capacity to incorporate improved management, this is not being directed towards species like Guitarfishes. When compared to requiem sharks (family Carcharhinidae), the average score for understanding stock status was less than half for Guitarfishes (18.3% vs. 7.1%), both of which are concerning scores (Sherman et al. 2022a). The pattern of attribute scoring of species was similar to that for countries, with greater scores for structural attributes and lower scores for fisheries operational attributes. However, the taxonomic resolution of landing limits scored marginally lower than species-specific compliance measures at the species level (14.8% ± 4.9 compared to 15.0% ± 3.3, respectively) (**Figure 4B**). The discrepancy between the score of the landing limit attribute across countries compared to species may be because some species with high scores are only found in a single country, weighting their scores higher when averaging country scores. Nineteen countries have no form of landing limits for any Guitarfish (68%) and 24 species (73%) have no landing limits of any kind in any country they are found in.

**Figure 4.**
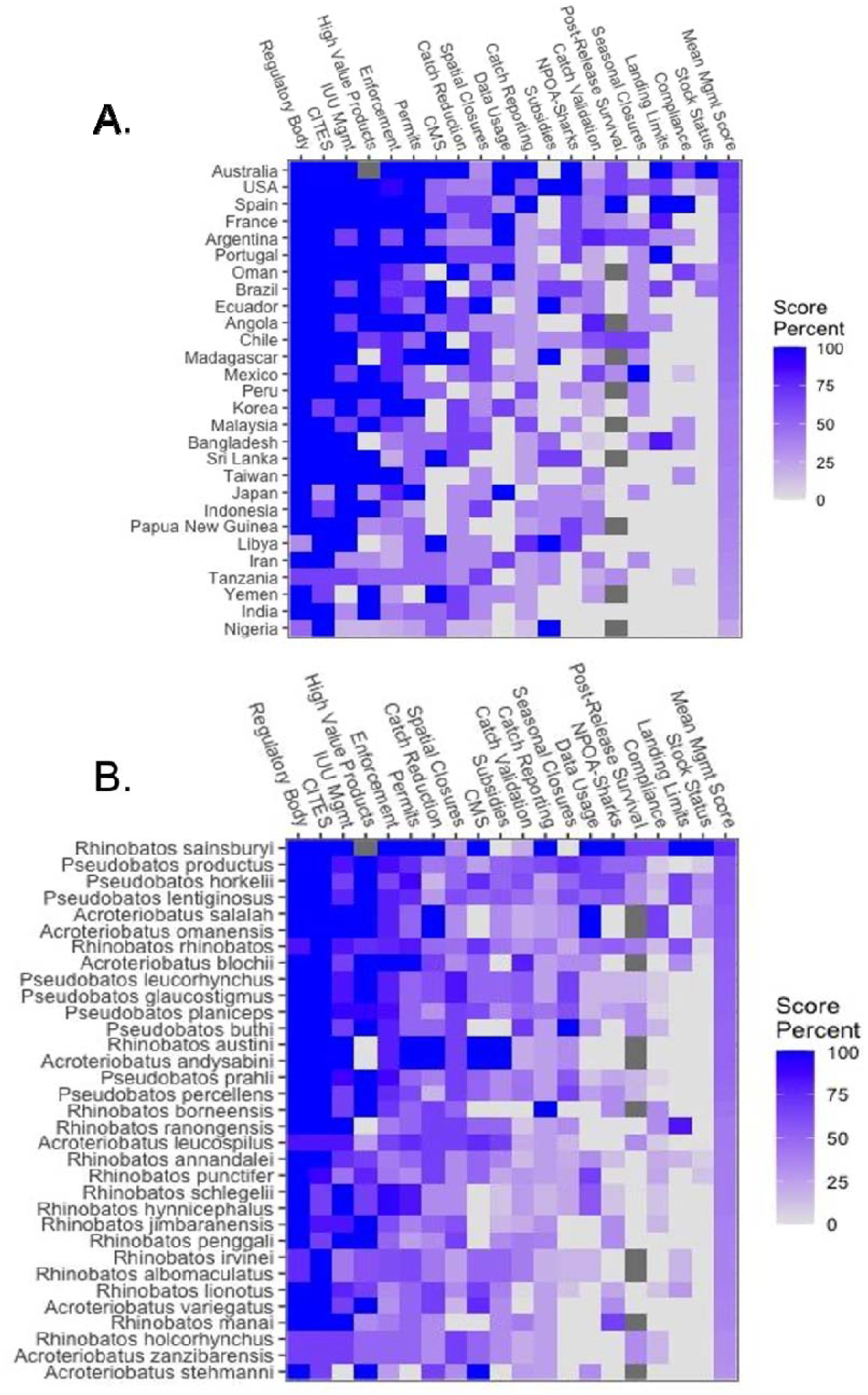
(*This figure should be colour). All average attribute scores and calculated management scores for all assessed species of guitarfish according to: (A) different countries, and (B) species. Attribute columns are arranged from highest average score to lowest average score. NA values are represented by dark grey squares.

## 4 Discussion

We find that, globally, there is likely to be a high volume of Guitarfishes present in international trade. However, their trade has been masked either by substantial underreporting, lack of studies, mislabelling, or non-specific commodity codes. Further, the management risk scores for all Guitarfish from 28 countries show they are undermanaged, with less than half of the ideal management in place (i.e., an average score of 45%). Globally, there are minimal species-specific regulations in fisheries, including a lack of landing limits, and catch being reported to a very aggregate taxonomic level, if reported at all. Due to the value of their fins, meat, snouts, and skins, there is well-documented international trade of Guitarfish products. Exports of Guitarfishes primarily originate from countries in Africa and Asia (Kyne et al. 2017, Jabado et al. 2021a, Jabado et al. 2021b), the two regions with the lowest management scores for these species. Consequently, the catch, trade, and undermanagement of Guitarfishes have resulted in an elevated extinction risk with over three quarters (77%) of Guitarfish species listed in a threatened IUCN category (i.e., VU, EN, CR) (IUCN Standards; Petitions Subcommittee 2019, Dulvy et al. 2021, IUCN 2022). Here, we consider the following: (1) the need to improve domestic catch regulations, (2) improving species-specific catch, landing, and trade data, (3) how management of Guitarfishes compares to other shark and ray taxa, and (4) the potential benefits of international trade regulation of Guitarfishes.

The most basic improvement in management that should be implemented at the national level, is increased species-specific reporting of Guitarfish catch. Reconstructed global catch of Guitarfishes shows boom and bust trajectories, with changes in which countries dominate the catches (Pauly et al. 2021). However, looking at global catch at the family level is misleading, particularly for species like Guitarfishes that have small geographic distributions. The shift in major Guitarfish-catching countries likely indicates population crashes in some countries, followed by increased prices and, therefore, demand filled by another country (with different species). In locations where there are many species, such as Ghana, serial depletions of adult and larger species of Guitarfishes are already being observed in regions where there is heavy fishing pressure and limited management (Seidu et al. 2022b). In other regions, such as the Bay of Bengal, fishers state there have been drastic declines in their Guitarfish catch throughout their time fishing, following the recent rapid collapse of Wedgefish and Giant Guitarfish populations (Haque et al. 2018). As the more high-value and high-demand species of Wedgefishes and Giant Guitarfishes are depleted, there is more incentive to mislabel products to achieve higher prices, further confusing the state of species’ populations (Alvarenga et al. 2021). Additionally, fishers are increasingly retaining ‘trash fish’, including species of Critically Endangered Guitarfishes, to increase their profits (Bhagyalekshmi and Kumar 2021). Therefore, implementing improved national management, including better species-specific catch reporting and retention limits may reduce the level of overexploitation and secure a more stable and longer-term future for local fishers.

Globally, there is a growing market for shark and ray meat (Dent and Clarke 2015, Liu et al. 2021). Due to the broad labelling of processed seafood to commodity codes, it is impossible to determine which species are included in this trade after processing has occurred (FAO 2022). The use of species-specific reporting of sharks and rays has increased in some countries following a species’ CITES listing in order to accommodate international trade (i.e., Indonesia (Prasetyo et al. 2021)). Currently, few countries report catch of Guitarfishes to FAO, including Brazil, which has not collected landing information since 2007 (Bernardo et al. 2020). Therefore, there is no data available to determine if catches (and by inference, populations) are increasing, stable or decreasing. Additionally, there is ongoing taxonomic discovery and revision of this family (i.e., (Weigmann et al. 2021)), further complicating our understanding of their inclusion in international trade. For example, despite many samples of the Rhinopristiformes order being found in markets in Hong Kong and Guangzhou, no Rhinobatidae were observed (Cardeñosa et al. 2020). This may be due to the unresolved taxonomy of Guitarfishes, which may result in misidentified or misclassified sequences in GenBank (Cardeñosa et al. 2020). Hence, the current poor-resolution of markers, some of which were developed prior to the major taxonomic reorganisation of rhino rays in 2016, mean that the occurrence of Guitarfishes cannot be ruled out with such molecular surveys of fin-clips. Finally, many Guitarfish products are imported and re-exported through several countries before reaching their final destination, which introduces traceability issues (Haque and Spaet 2021, Seidu et al. 2022b). In order to improve traceability and transparency, introducing international regulations that require exporters to prove the products were sustainably sourced and not detrimental to the population are needed.

Overall, Guitarfishes have lower management scores than requiem sharks, particularly in documented understanding of the stock status (Sherman et al. 2022a). Requiem sharks (Family: Carcharhinidae) also had low scores in the stock status attribute. This score is concerning because we have more information about requiem shark catch, trade, and high levels of threat (Cardeñosa et al. 2020, Dulvy et al. 2021, FAO 2022). However, despite the minimal understanding of requiem shark stock status, the average score for Guitarfishes was less than half that of requiem sharks, indicating how limited our understanding of their status is. Unlike requiem sharks, where widespread species were likely to have average management scores when compared to endemics (Sherman et al. 2022a), few Guitarfishes are widely distributed (**Figure 1a**). Many Guitarfishes are endemic to one or a few countries, therefore, we do not have assessments for all species. This also means that the management in each country a Guitarfish species occurs in has a much higher weighting or bearing upon the species’ global population and status. Whereas the more wide-ranging requiem sharks are likely to have a pocket of their range in a country with a better management risk score (i.e., USA or Australia), Guitarfishes that are endemic to undermanaged regions, particularly those with high levels of fishing such as Asia (Pauly et al. 2021), do not have these management ‘refuges’. Even still, the slightly more widespread species (i.e., common Guitarfish) are subjected to fragmented management, meaning their regional population trends may differ (Dulvy et al. 2017). Ensuring consistent management throughout their range may improve the overall species’ status. This can be done through international management plans or international agreements, like CITES.

Species listed on CITES Appendix II are subjected to international regulations regarding their trade. These regulations require national-level management to meet minimum standards, often requiring improvements, should a country wish to continue to export a listed species. This can be accomplished through Non-Detriment Findings (NDF), which require any traded specimens to be legally acquired sustainably (Fernando et al. 2022). Alternatively, some countries that are resource-poor may implement blanket bans on the landing of sharks and rays (Dulvy et al. 2017). We stress that this should not be the solution as blanket bans may have detrimental effects on subsistence and artisanal fishers who rely on these resources (Haque et al. 2022, Seidu et al. 2022a). Alternatively, they may force catches ‘underground,’ further complicating collection of landings data (Castellanos-Galindo et al. 2021). Therefore, we encourage national and regional level legislation with the goal of sustainable Guitarfish catch. This includes engaging with fishers to collect data and implement local policies that are co-designed with fisher input (Haque et al. 2022). Including fishers in the process may improve understanding of regulations and fisher compliance more efficiently than blanket bans alone (Booth et al. 2020). We consider that a CITES Appendix II listing of Guitarfishes would be a driver for change for nations where management is currently inadequate to ensure population recovery (where required) and achieve sustainable catches. In order to ensure long-term sustainability, both for Guitarfishes and the associated fisheries as food and income sources, improved management is essential.

### 4.1 Policy Relevance

There is a current proposal for Guitarfishes to be listed on the CITES Appendix II, which will be voted on in November of this year (2022). Our results show that most / almost all Guitarfishes have less than half (50%) of what would be considered ‘ideal management’ and over three quarters (77%) of data sufficient species are listed in a threatened category on the IUCN Red List (Dulvy et al. 2021). Despite the poor documentation for international trade of Guitarfishes, there is evidence from many countries that these species are traded and this may well be incentivising local fisheries and increasing extinction risk. As many of these species are predominantly found in developing countries with limited resources and capacity for management, it is likely their catch and trade is underreported, if reported at all. Overly broad commodity codes for international trade make it impossible to know the true volume of these species that are traded, as they would be listed under broad categories of fresh or processed fish/shark/ray. We conclude that listing Guitarfishes on CITES Appendix II provides a mechanism for tackling the management deficits for these species, incentivising national management, and increasing our understanding of their catch and trade. Further, listing Guitarfishes on Appendix II will harmonise the CITES listing of all rhino rays which can only ensure ease of implementation ensuring the recovery of this iconic evolutionary distinct lineage of fishes.

## Acknowledgements

The authors thank Cassandra L. Rigby for her comments on the manuscript. We also thank the following people for help finding or understanding legislation in various countries: Maite Pons (Argentina), David Kulka (Canada), Sushmita Mukherji and Zoya Tyabji (India), Aiko Matsushiba (Japan), Sara A.A. Al Mabruk (Libya), Gonzo Araujo and Cat McCann (Malaysia), Brittany Finucci (New Zealand), Leontine Baje and Michael Grant (Papua New Guinea), Andrew Bickell (Sri Lanka), Crystal McRae (Taiwan), and Tobey Curtis and John Carlson (USA).

