## Supplementary Information for "Guitarfishes are plucked: undermanaged in global fisheries despite declining populations and high volume of unreported international trade"

**Spreadsheets:**

**Data S1.** Countries with documented Guitarfish catch, export, and/or import, how catch is reported to the FAO where applicable, and sources used to determine catch and trade data.

**Data S2**. Average scores for each management unit including the scores of all species assessed in the management unit, SEM, number of countries each species was assessed in, number of species assessed in each country, and IUCN Red List Status of each species.

**Description of Attribute Classes Used for M-Risk**

**Management System**

These five attributes consider the foundational requirements for a sustainable fishery. including: (1) whether there is a regulatory body (i.e., fishery department), (2) the presence and structure of fishing permits, (3) the existence and depth of stock assessments for target species, (4) the existence and outcome of a risk assessment for the species being assessed, and (5) efforts to reduce shark catch (if not targeted).

***Fishing Practices & Catch***

These five attributes consider efforts to minimise fishing mortality through management of at-sea fishing operations. First, the landing limits attribute considers the taxonomic resolution of landing limits. Second, post-release survival (PRS) evaluates both the understanding of PRS for each species and regulations in place to increase PRS, thus reducing fishing mortality. Third, the high-value product attribute evaluates the effectiveness of finning regulations. The last two attributes in this group consider the presence and purpose (i.e., for target species, sharks overall, or species-specific) of spatial and temporal closures in-place.

**Compliance, Monitoring, & Enforcement**

This class of five attributes considers how fishing activity is examined, including: the taxonomic resolution of reported catch, how catch is validated, and any additional compliance measures not considered in other attributes. One attribute evaluates the consideration of illegal, unregulated, and unreported (IUU) fishing in management decisions. The last attribute considers enforcement methods including electronic surveillance, boarding of vessels, monitoring of catch unloading, logbook validation, and checking for illegal and mis-sized catch by fisheries officers.

**Country Attributes**

These four attributes indicate the importance of sustainable fisheries to a country’s government. Having a National Plan of Action for Sharks (Shark Plans) indicates a country’s interest in ensuring sustainable fishing mortality for sharks, with higher scores awarded for more thorough Shark Plans. Two country-level attributes consider engagement with international treaties and agreements, with higher scores awarded to countries that are CITES members with no reservations on sharks and higher scores for member countries of CMS and/or MOU-Sharks signatories. The final attribute in this class evaluated the national spending on fisheries subsidies and the proportion of deleterious “capacity-enhancing” subsidies compared to beneficial subsidies.

**Full Attribute Value Statements (from Sherman *et al., 2022*)**

**Management System**

***1.***	***Regulatory Body in Place***

Having a regulatory body allows fisheries to be managed. The regulatory body includes the views of fisheries managers, environmental groups, fishermen, and scientists, among others. This ensures that all invested parties have a say in the regulations made in the fishery. Without a regulatory body, a fishery is unable to adequately monitor catch. If this body does not meet regularly, changes in stock status or other stuff may not be addressed in a timely fashion. The score for this attribute will remain the same for all species assessed within a management unit.

0.	No regulatory body in place

1.	Regulatory body that meets less than once every two years

2.	Regulatory body that meets at least once every two years

3.	Regulatory body that meets at least once every year

***2.***	***Permits***

Having permits within a fishery enables managers to determine the number of fishers and allows for easy distribution of new legislation to those that are required to abide by it. Once permits are required, actual fishing pressure can be more accurately calculated. Having a permit that is associated with some form of fishing limit (ITQs or TAC) will enable fisheries managers to better control fishing mortality and adjust if overfishing is occurring. The score for this attribute will remain the same for all species assessed within a management unit.

0.	No permits required

1.	Permits / vessel registry required

2.	Permits with ITQ / TAC / trawl hours / some form of limit associated with the permit

***3.***	***Stock Status / Risk Assessments***

Collecting information regarding stock status is important to ensure overfishing is not occurring and that a population is not overfished. Once information is collected, stock status can be used to assess risk to overfishing. These assessments provide feedback to managers on the sustainability of the fishery and the ecosystem in which the fishery is operating. These assessments can expose areas of the fishery that require further research or can show that there is no need to complete more expensive assessments. In many cases, collating this information and performing an in-depth risk assessment is costly and time consuming, thus it is impractical. If basic risk assessments are performed and indicate there is no need for further assessments, stock status can be assumed to be healthy. In fisheries with the capacity to perform more in-depth stock assessments, results can be useful for smaller, adjacent fisheries. The score for this attribute may differ for each species in the management unit.

0.	No stock status / risk assessments performed

1.	Basic risk assessments performed showing high risk with no follow up assessments performed OR no provisions made for high risk species of elasmobranchs

2.	In depth assessments performed show overfishing is occurring OR basic assessment performed and some provisions made for high risk elasmobranchs

3.	In depth stock assessments show that overfishing is not occurring OR basic risk assessments performed showing no need for further assessments on the species (low risk fishery)

***4.***	***Sustainable Fishing Determination / Data Usage***

The method used for a fishery to determine the total allowable effort or catch (TAE / TAC) of their target species can inform what capacity the fishery has to manage its bycatch species. When determining TACC or total allowable effort, the method used will inform how accurate and safe the allowances are. With more data collected and available, more precise analyses can be done and, therefore, there is a better idea of a sustainable level of fishing mortality. This is measured for the target species that receives the lowest score, whether or not that is an elasmobranch. The method for determining sustainable fishing limits is telling of the overall management and resources available within the management unit. The score for this attribute will remain the same for all species assessed within a management unit.

0.	None

1.	Based on ERA results, estimations, or previous TACC/ effort

2.	Based on CPUE

3.	Based on stock assessments

***5.***	***Efforts to Reduce Catch***

In fisheries in which the species being assessed is not targeted, efforts to reduce bycatch should be in place to lower overall fishing mortality of the species. Methods to reduce catch of sharks and rays include bycatch reduction devices (BRDs) like turtle excluder devices, depth restrictions, trigger points, move-on limits. BRDs are frequently used in trawl fisheries and can significantly reduce the catch of larger elasmobranchs (Griffiths *et al.,* 2006). Depth restrictions have also been shown to reduce elasmobranch bycatch and in some instances, increase commercial value (Clarke *et al.*, 2015). Trigger points allow fisheries managers to identify a sustainable maximum catch and then stop commercial fishing operations once this has been hit for the year. Move-on limits are used in some fisheries and can be effective for species that aggregate. These limits require vessels to move a predetermined distance from the last point their gear was deployed if a certain level of bycatch is met to ensure another deployment does not completely remove the species in question from the area (Federal Fisheries Council, 2013; Nowara *et al.,* 2017). The score for this attribute may differ for each species in the management unit.

0.	No efforts in place to reduce catch

1.	One catch reduction method in place

2.	Two catch reduction methods in place OR one scientifically based catch reduction method

3.	Three or more catch reduction methods in place OR two or more scientifically based catch reduction methods

**Fishing Practices / Catch**

***6.***	***Taxonomic Resolution of Landing Limits***

Having science-based landing limits in place is the only direct method to impact fishing mortality of a species. Landing limits can be set based on previous catch, ERAs, scientific advice, or stock assessments, among other methods. Having landing limits for all sharks or rays is a first step in limiting the amount that can be taken. However, this limit may not be beneficial to some threatened species. For example, catching 10 tonnes of the Endangered Whale Shark (*Rhincodon typus*) would have a much bigger impact on the ecosystem than catching 10 tonnes of a Least Concern species, like the Milk Shark (*Rhizoprionodon acutus*). Species-specific catch limits are required to maintain a healthy elasmobranch community. The score for this attribute may differ for each species in the management unit.

0. No Landing limits set

1. Landing limit for CLASS / SUBCLASS “sharks / rays”

2. Landing limit for ORDER / FAMILY “hammerheads, thresher, deepwater sharks, etc.”

3. Species-specific landing limits OR no retention including the species being assessed

**7.**	***Removal of High Value Products (HVP)***

Many elasmobranchs have high value products (HVPs) that can be easily removed at sea. Vessel space is limited, thus fishers may remove HVPs at sea and discard the rest of the carcass in order to save valuable space and maximise their profitability. HVPs include fins, livers, *Mobula* gill plates, and rostra, among others. HVP at-sea removals have the potential to curb our ability to monitor species-specific catches. Fisheries have a few options to reduce HVP removal. These regulations ensure that entire carcasses are not wasted and when enforced, likely reduce cryptic mortality. The score for this attribute may differ for each species in the management unit.

0.	No regulations against removing any high value products at sea

1.	Fin to carcass ratios on landed sharks OR only regulations relating to removal of some HVPs relevant to the management unit

2.	Fin to meat ratios on landed sharks OR regulations relating to removal of all HVPs relevant to the management unit

3.	Fins naturally attached on landed sharks AND regulations relating to at-sea removal of all HVPs relevant to the management unit

***8.***	***Seasonal Closures***

Seasonal closures in fishery grounds allow for populations to naturally grow in the absence of fishing pressure. These closures are often implemented to allow for target species to spawn, breed, and/or pup. The reduction of fishing pressure can either incidentally or directly reduce fishing mortality of elasmobranchs, depending on the intention. In some areas, seasonal closures are in place, however, due to weather there would be no fishing operations at these times so they do not reduce fishing mortality further. The score for this attribute may differ for each species in the management unit.

0. No closures OR seasonal closure unlikely to curb fishing mortality (i.e. closed for monsoon season)

1. Seasonal closure for target species (non-elasmobranch) spawning / protection OR overall reduction in catch

2. Seasonal closure in place with the intention of reducing overall elasmobranch mortality

3. Seasonal closure for the benefit of the species being assessed (i.e. breeding/pupping grounds during season)

***9.***	***Spatial Closures***

Spatial closures can be put in place to protect important habitats for certain species, or a portion of a species’ range. These closures are often implemented in known sensitive locations for target species including areas for spawning, breeding, and/or pupping. The reduction of fishing pressure can either incidentally or directly reduce fishing mortality of elasmobranchs, depending on the intention. In some cases, spatial closures are put in place to benefit bycatch species, including some elasmobranchs. The score for this attribute may differ for each species in the management unit.

0.	No spatial closures

1.	Spatial closures in place for non-elasmobranchs

2.	Spatial closures in place for elasmobranchs but not the species being assessed

3.	Spatial closures in place for the species being assessed

***10.*** ***Post-Release Survival***

Post-release survival (PRS) estimates the likelihood of survival of animals caught by fishing gears and released back to the ocean. Estimates are based on condition of individuals as they are released. These estimates can be substantiated with tagging studies. Estimates are important because, when known, there can be measures put in place to increase the survival. Additionally, if there is high PRS, sustainability is less of a concern. As well as an estimate, measures can be taken to increase the likelihood of bycatch survival. These include circle hooks, which are more likely to hook a shark in the mouth than a deep hooking, and hoppers that reduce the time spent out of water while being sorted on the boat. Additionally, some fisheries include a code of practice on how to handle sharks in a manner that increases their survival once released. The score for this attribute may differ for each species in the management unit.

0.	No indication of PRS estimate or measures to increase PRS

1.	A post-release survival estimate

2.	Measures to increase post-release survival (through fishing gears modification i.e. circle hooks, hoppers, and or modification of handling practice e.g. Code of practice on shark handling to increase post-release survival)

3.	Both a PRS estimate and measures to increase PRS

**Compliance, Monitoring, and Enforcement**

***11. Catch Reporting***

Collecting catch information including ALL non-retained catch is important in understanding potential fishing mortality of a species. Species-specific information is best to fully understand the impact that a fishery has on a given species. As some species are more susceptible to post-release mortality, their catch levels may be unsustainable even without retention of the animals. For countries, when data cannot be found for the specific fishery, this will be scored by the most recent taxonomic level of reporting to FAO. Scores are only given if there are numbers associated with the species catch (i.e. a list of species caught in a fishery without the amount of each species caught is not considered). The score for this attribute may differ for each species in the management unit.

0.	No reporting of elasmobranch catch

1.	Catch reported to broad categories (i.e. “sharks” “elasmobranchs” or “rays”) OR list of species listed with no associated numbers of actual catch

2.	Catch reporting to narrow categories (i.e. “mackerel sharks,” “reef sharks,” “deepwater sharks,” “whaler sharks,” etc.)

3.	Catch reporting of similar/related species grouped (“deepwater dogfishes,” “gulper sharks,” “mako,” “hammerhead,” etc.)

4.	Species-specific catch reporting

***12.*** ***Illegal, Unregulated, and Unreported (IUU) Fishing***

Illegal, unregulated and unreported (IUU) fishing is hazardous as the complete fishing mortality of each species is not known. While IUU fishing cannot be stopped 100% due to the vast space of the ocean and relatively small size of the boats, it can be considered in management decisions including quota considerations and increased patrolling in areas with frequent IUU. The score for this attribute will remain the same for all species assessed within a management unit.

0.	IUU fishing a problem but not recognised in management documents

1.	IUU fishing a problem and acknowledged in management documents

2.	IUU fishing a problem and management arrangements reflect the impact IUU has in the management unit

3.	IUU fishing not a problem OR management arrangements to address IUU appear successful OR a signatory to the Agreement on Port State Measures (PSMA)

***13.*** ***Compliance Regime***

Fisheries management includes both generic and species-specific measures in order to control for fishing mortality. Some generic measures include limited entry (licensing system), gear restrictions, permanent area closures, etc. These regulations often will positively affect both target and non-target species populations by reducing fishing pressure and providing refuge areas. Species-specific measures include size limits, gender restrictions, move-on provisions, etc. These are commonly put in place for frequent bycatch species due to the impact fishing pressure may have on their populations. Not all of these regulations are relevant to the fishing mortality of each species assessed, therefore, they will be assessed based on whether the generic and/or species-specific measures are likely to reduce fishing mortality. The score for this attribute may differ for each species in the management unit.

0.	No species-specific measures in place / no relevant compliance measures in place

1.	Compliance measures in place unlikely to significantly reduce fishing mortality

2.	Compliance measures in place likely to reduce fishing mortality

3.	Species-specific measures in place that have proven to reduce fishing mortality

***14.*** ***Catch Validation***

All fisheries need some form of monitoring of fishing activities. Monitoring allows for managers to determine the level of activity and compliance within the fishery. Forms of monitoring include logbooks, various electronic monitoring systems, and observer programs or video observer programs (only when recordings of ALL catch of elasmobranchs are available). The score for this attribute will remain the same for all species assessed within a management unit.

* Observer program must be permanently in place to receive the points. Some fisheries put observers in place sporadically, therefore, only those fisheries that have observers every year will be considered. Fisheries with observer programs that do not operate each year will not receive any points for the program.

One point each for:

·	Self-reporting through logbooks

·	EMS (if there is video that covers 100% of the hauling of equipment)

·	Observer program* with less than 33% coverage (1 point)

·	Observer program* with up to 67% coverage (2 points)

·	Observer program* with 100% coverage (3 points)

***15.*** ***Enforcement Methods***

Methods of enforcement are measures taken that deter fishermen from breaking regulations. Enforcement penalties should provide higher incentive for fishermen to abide by regulations than they would have to fish illegally. Enforcement can include fines, criminal records, even jail-time, depending on the severity of the infraction. Fisheries have several ways of acquiring information that will lead to punishments. More ways to find those breaking rules will lead to more enforcement and better deter those performing illegal activities, therefore, this attribute is measured in the methods taken of finding those illegally fishing. The score for this attribute will remain the same for all species assessed within a management unit.

One point each for:

·	Use of electronic surveillance (e.g. drones, and/or planes, and/or VMS) to fine boats that are outside of regulations

·	Boarding of vessels to ensure compliance

·	Port checks for under/ oversized catch and/or illegal species

·	Logbook validation

·	Monitoring and/or weighing of catch unloading

**Country Indicators**

***16.*** ***Subsidies***

Subsidies are provided to fishermen to ensure they are profitable. Some subsidies ensure profitability by enhancing the capacity of fishermen to increase their catch, often at the expense of sustainability. These deleterious “capacity-enhancing” subsidies include fishery development, boat, market infrastructure, port, tax exemptions, access, and fuel subsidies. On the contrary, some subsidies enable fishermen to continue fishing sustainably without a loss of profit. Beneficial subsidies include those for fisheries management, research and development, and marine protected area implementation and management. The score for this attribute will remain the same for all species assessed within a management unit.

0.	Amount given for deleterious subsidies more than double the amount for beneficial subsidies

1.	Amount given for deleterious subsidies higher than for beneficial subsidies

2.	Amount given for beneficial subsidies higher than for deleterious subsidies

3.	Amount given for beneficial subsidies more than double the amount given for deleterious subsidies

***17.*** ***NPOA-Sharks and its Effectiveness***

Having an NPOA for sharks and rays shows a country is interested in the welfare of its shark populations. However, if certain NPOA objectives are not addressed, then it is unlikely to provide effective solutions for mitigating fishing mortality of sharks. Ten objectives were proposed by Davis and Worm (2013) as the minimum content to include. The score for this attribute will remain the same for all species assessed within a management unit.

0. 0-3 objectives addressed in NPOA

1. 4-6 objectives addressed in NPOA

2. 7-9 objectives addressed in NPOA

3. All 10 objectives addressed in NPOA

***18.*** ***CITES Member***

Is the country a member country of CITES? If so, do they have any reservations on listed chondrichthyan species? Being a member country of CITES indicates an overall interest in conservation of at-risk species. However, if there are reservations in place for species that should have additional management, it negates the impact of being a CITES member. The score for this attribute may differ for each species in the management unit.

0.	Not a member CITES

1.	CITES member with reservations on >50% of listed chondrichthyan species OR a reservation on the species being assessed

2.	CITES member with reservations on <50% of listed chondrichthyan species

3.	CITES member with no reservations on chondrichthyan species

***19.*** ***CMS Member***

Is the country a member country of CMS or a signatory on the MOU-Sharks? The score for this attribute will remain the same for all species assessed within a management unit.

0.	Not a member of CMS

1.	CMS member OR MOU-Sharks Signatory

2.	CMS member AND MOU-Sharks Signatory
